# SUMO conjugation regulates the activity of the Integrator complex

**DOI:** 10.1101/2022.03.07.483123

**Authors:** Laureano Bragado, Melina Magalnik, Pablo Mammi, Agustín Romero, Nicolás Gaioli, Berta Pozzi, Anabella Srebrow

## Abstract

RNA pol II transcribes snRNA genes in close proximity to Cajal bodies, subnuclear compartments that depend on the SUMO isopeptidase USPL1 for their assembly. We show here that overexpression of USPL1 alters snRNA 3’-end cleavage, a process carried out by the Integrator complex. Beyond its role in snRNA biogenesis, this complex is responsible for regulating the expression of different non-coding and coding transcripts. We validated several subunits of the complex as SUMO conjugation substrates, and found that the SUMOylation of INTS11 subunit is regulated by USPL1. We defined Lys 381, Lys 462 and Lys 475 as *bona fide* SUMO attachment sites within INTS11 and observed that SUMOylation of this protein is required for efficient Integrator activity. Moreover, while an INTS11 SUMOylation deficient mutant is still capable of interacting with INTS4 and INTS9, its interaction with other subunits of the complex is affected. This mutant also shows a more cytoplasmatic localization than the wild type protein. These findings point to a regulatory role of SUMO conjugation on Integrator activity and suggest the involvement of INTS11 SUMOylation in the assembly of the complex. Furthermore, this work adds the Integrator-dependent RNA processing to the growing list of cellular processes regulated by SUMO conjugation.

## INTRODUCTION

In eukaryotic cells, protein-coding genes are transcribed by RNA polymerase II (RNAPII) generating pre-mRNAs that contain exons and introns. The splicing process removes introns and joins exons giving rise to mature mRNAs, and is catalyzed by the spliceosome. This highly complex and dynamic macromolecular machine is composed of five ribonucleo-protein particles (RNPs) termed small nuclear RNPs (snRNPs) U1, U2, U4/U6 and U5, in the case of the so called “major spliceosome”. Each snRNP comprises one small nuclear RNA (snRNA) associated with a specific set of proteins. While the function of snRNAs and the assembly of snRNPs along the splicing process are well characterized (1), the regulation of snRNA gene expression is still poorly understood.

Most snRNAs are transcribed by RNAPII, including U1, U2, U4 and U5. Unlike pre-mRNAs, snRNAs are intronless and non-polyadenylated. 3’ end processing of the nascent snRNA transcript depends on a sequence termed the 3’ box (2), which is recognized by the Integrator complex (3). In the absence of this complex, nascent snRNAs are cleaved and polyadenylated by the cleavage and polyadenylation machinery (CPA) that operates on pre-mRNAs (4).

The Integrator complex contains 14 subunits (INTS1-INTS14), where INTS9 and INTS11 have been identified as homologs of cleavage and polyadenylation specificity factors (CPSF) 100 and 73, respectively (3), members of CPA. Although INTS11 is responsible for the endonucleolytic activity, it has been shown that INTS4, INTS9 and INTS11 associate into a ternary complex, which constitutes the Integrator catalytic core. Furthermore, the structure of this core resembles that of the heterotrimeric complex CPSF73/100/Symplekin (5,6).

In recent years, several reports have linked Integrator complex activity with the biogenesis of a broader set of substrates, including enhacer RNAs (eRNAs), viral microRNAs and diverse non-polyadenylated long-non coding (lnc) RNAs (7–9,14–16). In addition, this complex is involved in the fine regulation of promoter-proximal pausing of RNAPII on protein coding genes (10–14,18– 20). Moreover, the Integrator complex is necessary for the transcriptional response downstream of growth factor signaling and INTS11 is phosphorylated by ERK1/2 (17). However, the relevance of post-translational modifications on the activity of INTS11, or other subunits of the Integrator complex, has not yet been addressed.

SUMO conjugation (aka SUMOylation) is a rapid, reversible post-translational modification (PTM) consisting in the covalent attachment of a small ubiquitin-related modifier (SUMO) peptide to a lysine residue in the target protein. There are three well-characterized functional SUMO isoforms encoded by the human genome (SUMO1, 2 and 3), which modify distinct but overlapping sets of substrates. While it is still unclear whether SUMO4 is conjugated to cellular proteins, SUMO5 has been more recently identified as a novel, primate-and tissue-specific SUMO variant (21–25). Like ubiquitin, SUMO is conjugated to its targets by an isopeptide bond between its C-terminal glycine and the ε-NH2 group of the target lysine residue. In general, SUMOylation substrates contain the consensus motif ΨKxD/E, where Ψ is a large, hydrophobic amino acid, K is the target lysine, x is any amino acid, D is aspartic acid and E glutamic acid. However, many SUMOylated proteins deviate from this consensus sequence or even lack one (26,27). The steps of the SUMO conjugation pathway resemble those of the ubiquitin pathway. Before being conjugated, SUMO is cleaved by specific proteases (SENPs), exposing its C-terminal Gly-Gly motif (28). Subsequently, mature SUMO is activated by the SUMO-activating enzyme E1, the heterodimer ‘AOS1-UBA2’, in an ATP-dependent manner and then transferred to the catalytic Cys residue of ‘Ubc9’, the SUMO-specific E2 conjugating enzyme. Finally, an isopeptidic bond is formed between the C-terminal Gly residue of SUMO and a Lys residue in the target protein. This step is generally aided by SUMO E3 ligases and among those characterized so far, few display substrate specificity while others display SUMO isoform preferences (22). SUMOylated proteins are substrates of SENPs and other isopeptidases, which deconjugate SUMO, ensuring the reversibility and dynamic nature of the process. Most frequently, SUMO conjugation regulates intra-or intermolecular interactions, altering either the conformation of the modified protein or the recruitment of its partners (22). In several cases, SUMOylation fosters new associations by non-covalent interaction of conjugated SUMO with proteins harboring SUMO-interaction motifs (SIMs). The establishment of SUMO–SIM interactions exerts a variety of effects, ranging from intramolecular structural rearrangements, as reported for thymine DNA glycosylase, to the assembly and stabilization of multi-protein complexes, as described for PML nuclear bodies (29). In addition, SUMOylation can also interfere with protein stability by triggering ubiquitylation of poly-SUMO-modified proteins through the recruitment of SUMO-targeted ubiquitin ligases (STUbL) (30).

The biological relevance of protein SUMOylation is clearly demonstrated by the fact that inactivation of SUMO in *Saccharomyces cerevisiae* or of the unique E2 SUMO-conjugating enzyme Ubc9 in mice is lethal (31,32). Consistent with this, multiple studies have shown that SUMOylation regulates a wide range of cellular functions, including intracellular transport, maintenance of genome integrity, formation of nuclear subdomains (21), and also some aspects of rRNA or snoRNA metabolism (33–35). Furthermore, SUMO conjugation affects not only the stability, localization, and activity of transcriptional regulators, but also the activity of DNA and histone modifiers, leading to changes in chromatin structure and hence gene expression (36).

Proteomic approaches have revealed that RNA-related proteins are predominant among SUMO substrates (37). In addition, Ubc9 has been found to localize in nuclear speckles (38), which are thought to coordinate splicing and gene expression, as they contain not only splicing factors, but also other proteins involved in mRNA metabolism, such as transcription factors, RNA polymerase II subunits, cleavage and polyadenylation factors, and RNA export proteins (39). Furthermore, SUMO conjugation regulates different aspects of mRNA metabolism, such as pre-mRNA splicing (40), pre-mRNA 3’ end processing and RNA editing, by modifying the function of prp3 spliceosomal protein, poly(A) polymerase, Symplekin and CPSF-73, and ADAR1 respectively (41,42).

RNAPII transcribes snRNA genes in close proximity to Cajal bodies (CBs) (43–47) and coilin, a protein known to function as a scaffold for CBs, has been shown to directly interact with snRNAs (48). The Little Elongation Complex (LEC), necessary for elongation of RNAPII during transcription of snRNAs, colocalizes with coilin at CBs (49). In addition, the SUMO protease USPL1 has been found to localize with coilin within these nuclear bodies and to interact with the LEC. Although USPL1 knockdown causes diminution of CB formation and a reduction of snRNA levels, the relevant substrates of USPL1 have not been identified and the molecular mechanism underlying these effects are still unclear (50,51).

In this study, we have investigated the possible involvement of SUMO conjugation in snRNA biogenesis. We verified several subunits of the Integrator complex as *bona fide* SUMO conjugation substrates in cultured human cells. In particular, we focused on the catalytic subunit of this complex, INTS11, mapping the target residues for this modification and demonstrating that its SUMOylation levels are regulated by the SUMO isopeptidase USLP1. In addition, we have shown that USLP1 affects snRNA 3’end processing. By generating a SUMOylation-deficient mutant of INTS11 we found that SUMO conjugation to this subunit is not only crucial for proper snRNA biogenesis but also for the expression of other transcripts including certain eRNAs and prompts, two classes of non-coding RNAs recently found to be regulated by the Integrator complex. We propose that while the assembly of the catalytic core is not dependent on INTS11 SUMOylation, this modification is required for the proper sub-cellular localization of this subunit as well as for its interaction with components of the Integrator complex other than INTS4 and INTS9. These results reveal a novel regulatory mechanism for Integrator activity and add Integrator-dependent RNA processing to the list of cellular processes regulated by SUMO conjugation.

## MATERIALS AND METHODS

### Cell lines

HeLa (human cervix adenocarcinoma cell line, ATCC CCL-2) and HEK 293T cells (human embryonic kidney cell line, ATCC CRL-1573) were maintained in Dulbecco’s Modified Eagle’s Medium supplemented with 10% fetal bovine serum, 100 U/ml penicillin and 100 μg/ml streptomycin.

### DNA plasmids

Expression vector for FLAG-INTS11 was a generous gift from Dr. Ramin Shiekhattar (University of Miami Health Systems, US), expression vectors for FLAG-INTS9, FLAG-INTS10, HIS-MYC-INTS9, HIS-MYC-INTS11 were a generous gift from Dr. Shona Murphy (University of Oxford, UK), expression vectors for HA-INTS4, HA-INTS13, HA-INTS14 were a generous gift from Dr. Stefanie Jonas (Institute of Molecular Biology and Biophysics, ETH Zurich), expression vector for HA-USPL1 was a generous gift from Dr. Gideon Dreyfuss (University of Pennsylvania, US) and U7 snRNA-GFP reporter was a generous gift from Dr. Omar Abdel-Wahab (Memorial Sloan Kettering Cancer Center, NY, US).

The expression vector for GFP-FLAG-INTS11 was generated in our laboratory. Briefly, INTS11 cDNA was amplified from FLAG-INTS11 plasmid with the following primers: Forward: AAAGAATTCTGACTACAAAGACGAT and Reverse: AAAGGATCCTCTAGAGTCGACTGGT. PCR product was digested with EcoRI and BamHI restriction enzymes and sub-cloned into EcoRI/BamHI-digested pEGFP-C1 plasmid (Addgene #2487).

### Transfection

Transfection of plasmid DNA and siRNA was carried out with Lipofectamine 2000 (Thermo Fisher) according to manufacturer’s instructions.

### Western blot assay and antibodies

Protein samples were resolved by SDS-PAGE and transferred to nitrocellulose membranes (BioRad). Membranes were blocked and then incubated with the corresponding primary antibody. After washing, membranes were incubated with IRDye® 800CW (LI-COR Biosciences) secondary antibody. Bound antibody was detected using an Odyssey imaging system (LI-COR Biosciences). Western blots were performed at least three times, and representative images are shown in each case. The antibodies used were: mouse monoclonal anti-β-actin C4 (Santa Cruz Biotechnology), polyclonal rabbit anti β-tubulin H-235 (Santa Cruz Biotechnology), monoclonal mouse anti GFP B2 (Santa Cruz Biotechnology), monoclonal mouse anti c-Myc 9E10 (Santa Cruz Biotechnology), monoclonal mouse anti FLAG-M2 (Sigma-Aldrich), monoclonal mouse anti HA MMS-101R (Covance), polyclonal rabbit anti INTS11 ab75276 (Abcam), polyclonal rabbit anti INTS11 A301-274A (Bethyl), polyclonal rabbit anti INTS9 13945 (Cell Signaling Technology)

### Site-directed mutagenesis

Mutagenesis was performed by the DpnI method, based on Stratagene’s QuickChange protocol. The primers used to mutate putative SUMO sites from Lys to Arg are listed in Supplementary Table S2. Mutations were always verified by sequencing.

### Purification of HIS-SUMO conjugated proteins

HEK 293T cells were transfected in 35-mm culture wells with the indicated plasmids. After 48 h, 6xHIS-SUMO2 conjugates were purified under denaturing conditions using Ni-NTA-agarose beads according to manufacturer’s instructions (Qiagen). Briefly, transfected cells were harvested in ice-cold PBS plus 100 mM iodoacetamide (IAA). An aliquot was taken as input and the remaining cells were lysed in 6M guanidinium-HCl containing 100 mM Na2HPO4/NaH2PO4, 10 mM Tris–HCl pH 8.0, 5 mM imidazole and 10 mM IAA. Samples were sonicated to reduce the viscosity and centrifuged for 20 min at 12000 g. Afterwards, proteins in the supernatants were purified using Ni-NTA beads (Qiagen) according to (62). Samples were subsequently washed with wash buffer I (8 M urea, 10 mM Tris-HCl, 100 mMNa2HPO4/NaH2PO4, 5 mM imidazole, 10 mM IAA, pH8.0), wash buffer II (8 M urea, 10 mM Tris-HCl, 100 mM Na2HPO4/NaH2PO4, 0.2% Triton X-100, 5 mM imidazole, 10 mM IAA, pH 6.3), and wash buffer III (8 M urea, 10 mM Tris-HCl, 100 mM Na2HPO4/NaH2PO4, 0.1% Triton X-100, 5 mM imidazole). Samples were eluted in 2X Laemmli sample buffer containing 300 mM imidazole (pH 6.3) for 3min at 95° C.

### Quantitative PCR for cellular RNAs (RT-qPCR)

Total cellular RNA was isolated by using 250 µl of Tri-Reagent (MRC) and measured with a NanoDrop 1000 spectrophotometer (Thermo Scientific). RNA was treated with DNase RQ1 (Promega) following the manufacturer’s instructions. Then, 1 µg of each RNA sample was reverse transcribed to cDNA with random deca-oligonucleotide or Oligo-dt primer mix using MMLV Reverse Transcriptase (Invitrogen). Quantitative PCRs (qPCRs) were performed using SYBR Green dye, 1/20 dilution of cDNA sample and Taq DNA polymerase (Invitrogen) in a Mastercycler® ep realplex PCR device (Eppendorf). The annealing temperature was 60 °C and the elongation time at 72 °C was 30 sec. Relative RNA abundance from cDNA samples and no-reverse transcription controls was estimated employing internal standard curves with a PCR efficiency of 100 ± 10% for each set of primers, in each experiment. Realplex qPCR software was used to analyze the data. The specific primers used are listed in Supplementary Table S2.

### CRISPRi-mediated depletion

For CRISPRi, 50,000 HEK 293T cells were seeded in 24-well plates. After 24 h, 400 ng of dCas9-KRAB expression vector (Addgene #60954), 170 ng of lentiGuide puro (Addgene #52963) and 30 ng of expression vector for INTS11 were transfected. Cells were harvested 72 h post-transfection. Sequences of sgRNAs are listed in supplementary Table S2

### Chromatin immunoprecipitation

Cells were cross-linked with 1% (v/v) formaldehyde (final concentration), washed twice with cold PBS, scraped, collected and centrifuged. Cell pellets were resuspended in 2 ml of SDS lysis buffer (1% w/v SDS, 10 mM EDTA, 50 mM Tris–HCl, pH 8.1) containing Complete Protease Inhibitor Cocktail (Roche) and incubated for 10 min on ice. Cell extracts were sonicated with a Branson sonicator W-450 D at 30% amplitude with fifteen 10-s bursts, resulting in ∼500-nt chromatin fragments and then centrifuged for 10 min at 12 000 g. A 50 µl sample of the supernatant was saved as input DNA and the remainder was diluted 1:10 in ChIP dilution buffer (0.01% w/v SDS, 1.1% v/v Triton X-100, 1.2 mM EDTA, 16.7 mMTris–HCl pH 8.1, 167 mM NaCl) containing protease inhibitors. The chromatin solution was precleared at 4°C with Protein G Dynabeads® for 1 h before incubating overnight at 4°C with antibodies. Complexes were incubated with Dynabeads® Protein G beads for 1 h at 4°C. Beads were washed by rocking for 4 min, once in each of the following buffers: low salt immune complex wash buffer (0.1%w/v SDS, 1%v/vTritonX-100, 2mMEDTA, 20mM Tris–HCl, pH 8.1, 150 mM NaCl), high-salt immune complex wash buffer (same as low salt buffer, except with 500 mM NaCl) and LiCl immune complex wash buffer (0.25 M LiCl, 1% v/v NP-40, 1% w/v deoxycholic acid, 1 mM EDTA, 10 mM Tris–HCl, pH 8.1), and then twice in TE (10 mM Tris–HCl, 1 mM EDTA). Bound complexes were eluted in 1% (w/v) SDS and 50 mM NaHCO3 and crosslinking was reversed by incubating for overnight at 65°C. Samples were digested with proteinase K for 1 h at 45°C and DNA extracted using a Qiagen PCR purification kit. DNA retrieved by ChIP was analyzed by quantitative real-time PCR with primers listed in supplementary Table S2. Data sets were normalized to ChIP input values.

### Immunoprecipitation of the Integrator complex

Cells were harvested and lysed in RIPA buffer (50 mM Tris-HCl pH 7.5, 1% (v/v) NP-40, 0.5% (w/v) sodium deoxycholate, 0.05% (w/v) SDS, 1 mM EDTA, 150 mM NaCl) containing Complete Protease Inhibitor (Roche). Extracts were sonicated at high amplitude with three 10-s bursts, and insoluble material was pelleted. Anti-FLAG M2 were added to the supernatant and incubated overnight. Then, complexes were incubated with Protein G Dynabeads for 1 h and washed three times in wash buffer (50 mM Tris–HCl (pH 7.5), 0.1% (v/v) NP-40, 1 mM EDTA, 125 mM NaCl). For western blot analysis, beads were resuspended in 2X Laemmli sample buffer. For GFP and GFP-INTS11 immunoprecipitations, cells were lysed in RIPA buffer (same as above) containing Complete Protease Inhibitor (Roche). Extracts were sonicated at high amplitude with three 10-s bursts, and insoluble material was pelleted. Supernatants were incubated 1h with GFP-Trap coupled to magnetic agarose beads (Chromotek). Then, complexes were washed three times in wash buffer (same as above). For western blot analysis, beads were resuspended in 2X Laemmli sample buffer.

### Microscopy confocal assays

HeLa cells were seeded into 24-well plates containing glass coverslips. Twenty-four hours later, cells were transfected with the indicated plasmids. After 24 h, coverslips were collected, and cells were fixed with paraformaldehyde (PFA) 4% in PBS pH 7.4 at room temperature for 10 min. PFA-fixed cells were then permeabilized with 1% Triton X-100 for 5 min at room temperature. Coverslips were incubated with RNASE A for 30 min at 37ºC. After three washes with PBS, incubation with propidium iodide was performed for 5 min and washed 4 times with PBS. Coverslips were mounted with a drop of mounting medium (Vectashield) and images were obtained with an Olympus FV1000 confocal microscope. Fluorescence intensity analysis of images was performed with Cell Profiler software (v 3.1.5) after generating a suitable pipeline for this purpose.

### Statistics

Typically, three or four independent experiments in triplicate repeats were conducted for each condition examined. Average values are shown with standard deviation and p-values, measured with a paired two-tailed t-test or ANOVA (as indicated for each case). Significant p-values are indicated by the asterisks above the graphs (p<0.001=***; p<0.01 = **; p<0.05 = *).

## RESULTS

### Overexpression of USPL1 affects the levels of nascent snRNAs

USPL1 is an essential SUMO isopeptidase that localizes to CBs and its knockdown leads to disruption of these sub-nuclear structures (50). Moreover, USPL1 has been shown to interact with components of the RNAPII-associated LEC and knockdown of USPL1 leads to reduced RNAPII-mediated snRNA gene transcription, diminished production of snRNPs and altered pre-mRNA splicing (51). However, USPL1 substrates implicated in these effects have not yet been identified.

To further study the involvement of USPL1 in snRNA biogenesis, we over-expressed HA-USPL1 together with a U7-GFP readthrough reporter construct (52) in human cultured cells (Figure 1A). Under basal conditions, marginal EGFP expression from the reporter is observed due to 3’-end cleavage of U7 snRNA gene transcripts, which impairs expression of the downstream EGFP ORF. However, when snRNA 3’-end processing is perturbed, e.g. by tampering with the expression of Integrator complex subunits, RNAPII transcribes beyond the U7 cleavage site (3’ box) and recognizes a canonical cleavage and polyadenylation signal (pA) downstream of the EGFP ORF (Figure 1A) (52). Over-expression of HA-USPL1 markedly increases GFP expression levels (Figure 1B and 1C) while partially decreasing global SUMOylation (Figure 1D), compared to the levels observed upon co-transfection of the reporter with empty plasmid (pcDNA). We then analyzed total and uncleaved levels of endogenous, RNAPII-dependent snRNA transcripts, by RT-qPCR (Figure 1E). The levels of uncleaved U1, U2, U4 and U5 snRNAs are increased upon over-expression of USPL1 (Figure 1F), suggesting that SUMOylation is involved in snRNA maturation.

**Figure 1.**
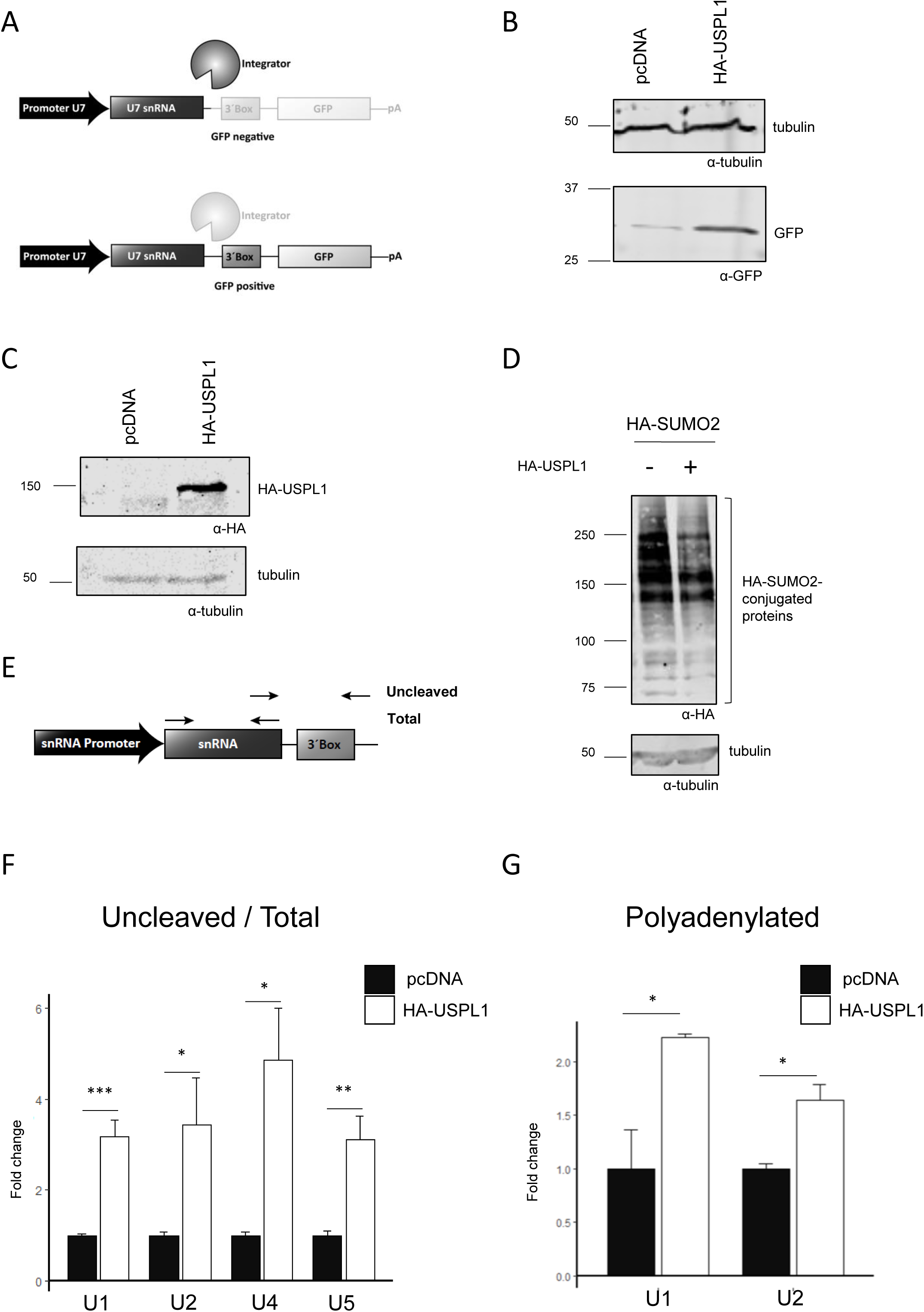
Over-expression of USPL1 affects nascent snRNA levels. (A) Schematic of U7-GFP reporter construct that produces Green Fluorescence Protein upon abrogation of Integrator complex activity. (B-C) HEK 293T cells were co-transfected with the U7-GFP reporter mentioned in A and HA-USPL1 expression vector. (D) Same as B and C, with the addition of HA-SUMO2 expression vector. In every case, after 48 h, whole cell lysates were subjected to western blot with the antibodies indicated below each panel. (E) Schematic of human snRNA genes showing the position of the primers used to amplify total and uncleaved snRNAs by RT–qPCR. (F-G) RT-qPCR analyses of RNA samples corresponding to cell culture conditions indicated in B-D, using primer pairs targeting the snRNA genes indicated below each graph. Reverse transcription was performed with random decamer primer (F) or oligo dT primer (G). Uncleaved snRNA levels were normalized to total snRNAs (F) or GAPDH mRNA (G). Average values are shown with standard error and P-values, determined using a paired two-tailed t-test. Significant P-values are indicated by the asterisks above the graphs (n=4, ***P< 0.001; **P< 0.01; *P< 0.05).

Previous studies have shown that in the absence of proper snRNA 3’-end cleavage, snRNAs become polyadenylated (4,9). Thus, the levels of polyadenylated snRNAs can be considered an indicative of 3’-end processing failure. Consistent with our above-mentioned observations, over-expression of HA-USPL1 also leads to increased levels of polyadenylated snRNAs (Figure 1G). These results demonstrate that maintenance of USPL1 levels is critical for efficient snRNA 3’-end processing. Whether this is due to a direct effect of USPL1 on components of the 3’-end processing machinery or to an overload of this machinery by an increase in snRNA transcription remains to be elucidated.

### Integrator subunits are modified by SUMO in cultured cells

Having observed that over-expression of a SUMO isopeptidase alters snRNA biogenesis, we analyzed human SUMO proteome data sets to further explore the possible effect of SUMO conjugation on the activity of snRNA 3’-end processing factors. Different proteomic studies revealed several Integrator complex subunits as SUMO2/3 conjugation targets (Supplementary Table S1) (27).

To validate their modification by SUMO conjugation in our culture conditions, cell extracts from HEK 293T co-transfected with expression vectors for 6xHis-SUMO2 and tagged Integrator subunits were subjected to nickel affinity purification, to enrich for SUMOylated proteins. Pulled-down proteins were analyzed by western blot with specific antibodies against the different fused tags. As shown in Figure 2, we were able to detect SUMO conjugation to HA-INTS4 (Figure 2A), FLAG-INTS9 (Figure 2B), FLAG-INTS10 (Figure 2C), FLAG-INTS11 (Figure 2D), HA-INTS13 (Figure 2E), HA-INTS14 (Figure 2F). It has been proposed that INTS4, INTS9 and INTS11 form the Integrator “cleavage module” responsible for 3’-end processing of target RNAs. INTS11 contains the active site that is responsible for Integrator activity, but its association with INTS4 and INTS9 is crucial for catalysis (5). After identifying that the three subunits of the Integrator cleavage module are *bona fide* SUMOylation substrates, we investigated the possible regulation of this modification by USPL1. To do so, we performed a nickel affinity purification strategy followed by western blot, as indicated above, but with total lysates derived from cells also over-expressing HA-USPL1. Remarkably, the levels of INTS11 SUMOylation, but not those of INTS4 and INTS9, are modulated by USPL1 (Figure 2G, 2H, and 2I), consistent with the above suggested involvement of SUMO conjugation in snRNA maturation.

**Figure 2.**
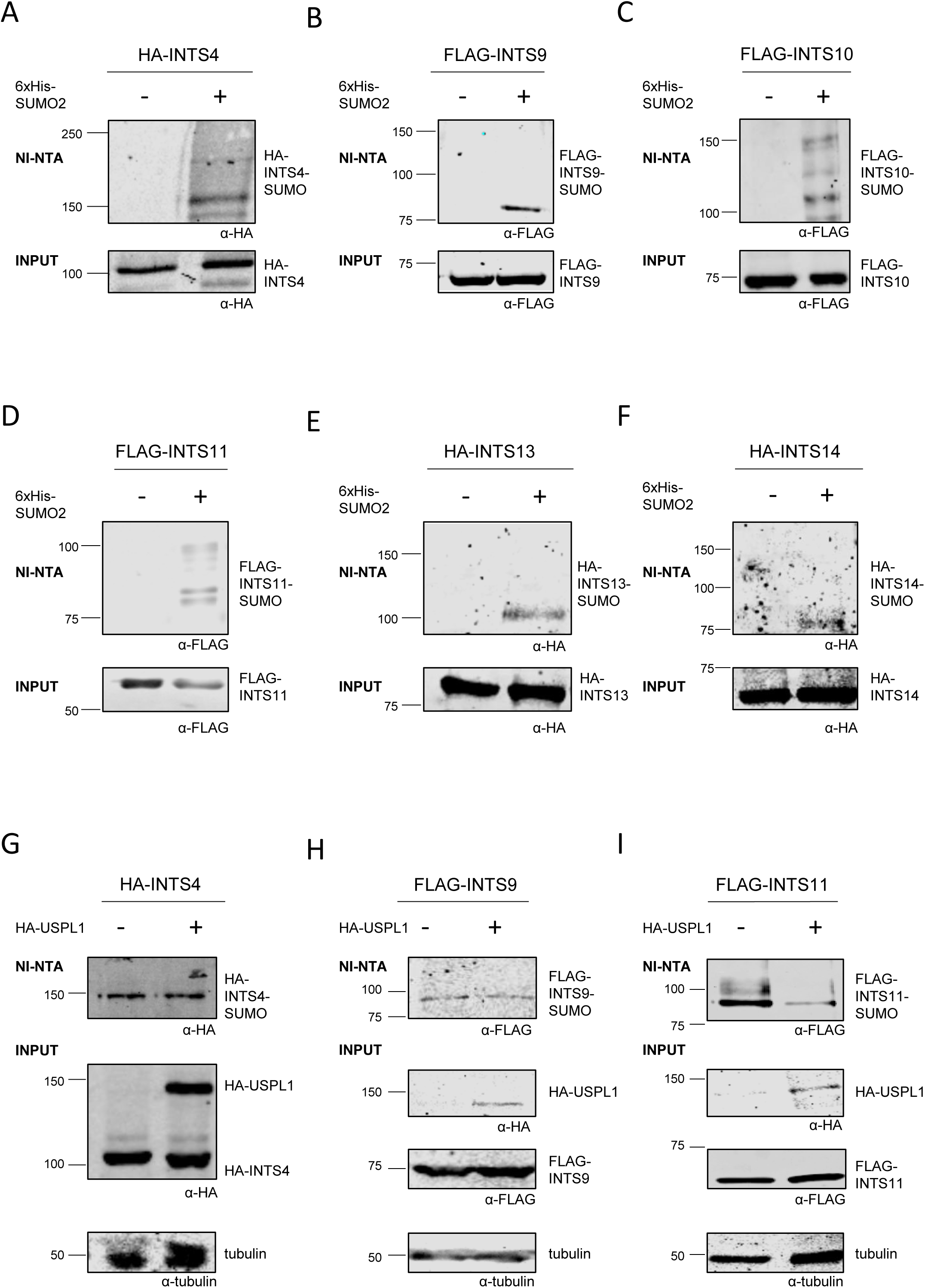
Integrator subunits are modified by SUMO conjugation in human cultured cells. (A-I) HEK 293T cells were transfected with expression vectors indicated at the top of each panel. After 48 h, cells were lysed and whole cell lysates were subjected to nickel affinity chromatography (Ni-NTA). Aliquots of the cell lysates (input) and eluates (Ni-NTA) were analyzed by western blot with the antibodies indicated below each panel.

### INTS11 is SUMOylated at lysines 381, 462 and 475

The cleavage module of the Integrator complex is closely related to the cleavage module of the CPSF complex that is required for pre-mRNA cleavage at poly(A) sites. Previous studies have not only shown that SUMO conjugation modulates pre-mRNA 3’-end processing but also that CPSF73, the subunit of the CPSF complex that catalyzes the endonucleolytic cleavage, is modified by SUMO. However, the consequences of this modification on CPSF73 catalytic activity are not completely understood (41). Taking into account that INTS11 shares sequence identity with CPSF73 (3), and that USPL1 regulates the levels of INTS11 SUMO conjugation, we focused on INTS11 SUMOylation.

Initially, bioinformatics prediction was used to define potential canonical SUMO acceptor sites within this protein (Sumo sp 2.0 <www.sumosp.biocuckoo.org>). This search revealed a putative SUMO conjugation site at Lys 462 of human INTS11, within the ΨKxD/E SUMOylation consensus motif. Site-directed mutagenesis was performed in order to replace this Lys by Arg, a widely-used strategy to map SUMOylation target residues. SUMOylation levels of the FLAG-INTS11 mutant was analyzed by western blot after nickel-affinity purification of SUMOylated proteins from cells co-expressing 6xHis-SUMO2 and the different FLAG-INTS11 versions. Mutation of Lys 462 provoked only a slight decrease in INTS11 SUMO conjugation levels, compared to the wild type version of this protein (Figure 3A). In addition to this canonical SUMO consensus sequence, we also identified a Lys residue within an inverted consensus motif D/ExKΨ, Lys 381. A double mutant was then generated, and the combined replacement of Lys 381 and 462 by Arg residues led to a marked reduction in INTS11 SUMOylation levels (Figure 3B). Site-specific SUMO proteomics data have identified Lys 115, 289, 369, 462 and 475 as high score SUMO acceptor sites within INTS11 (27). Based on this information, several INTS11 triple mutants were generated in order to further reduce INTS11 SUMOylation. Only the combined replacement of Lys 381, 462 and 475 by Arg residues (here after referred as “INTS11 3KR” mutant) led to a robust and drastic reduction of SUMO conjugation to INTS11 (Figure 3C and 3D). It is worth noticing that neither the FLAG-INTS11 catalytic-inactive mutant (E203Q) nor the GFP-FLAG-INTS11 fusion protein display any perturbation of their SUMOylation levels (Supplementary Figure S1A and S1B). The three Lys residues, and the SUMO consensus motif, are highly conserved across different species suggesting that SUMO conjugation of these proteins may also be conserved (Supplementary Figure S1C).

**Figure 3.**
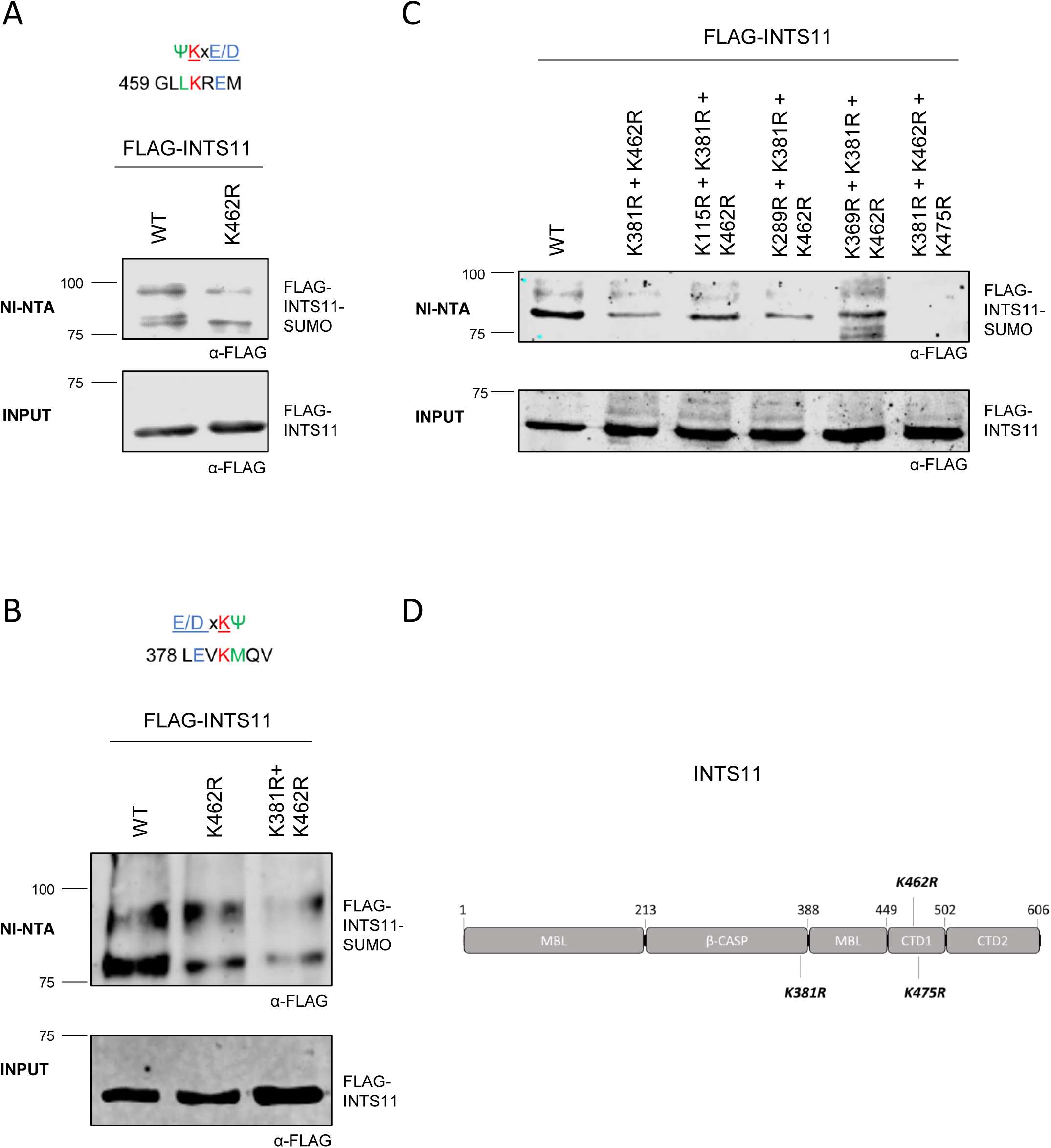
INTS11 is SUMOylated at lysine residues 381, 462 and 475. (A-C) HEK 293T cells were co-transfected with expression vectors encoding WT or mutated FLAG-INTS11, and 6xHIS-SUMO2, as indicated above each lane. After 48 h, cell lysates were subjected to nickel affinity chromatography (Ni-NTA). Aliquots of the cell lysates (Input) and eluates (Ni-NTA) were analyzed by western blot with an anti-FLAG antibody to detect the FLAG-INTS11 variants. (A-B) Upper panels show putative SUMO attachment sites in INTS11, predicted in silico, and the surrounding SUMO consensus motif. Ψ bulky, hydrophobic amino acid; X, any amino acid; E, glutamic acid; D, aspartic acid. (D) Domain architecture of INTS11 and localization of detected SUMO conjugation sites (bold).

Considering that the triple mutant “INTS11 3KR” showed the lowest level of INTS11 SUMOylation in our experimental setting, it was used to further analyse the consequences of SUMO conjugation on different INST11-mediated processes.

### Lack of INTS11 SUMOylation affects Integrator activity in cultured cells

To explore whether INTS11 SUMOylation has indeed any relevance for Integrator activity within cultured cells, we analyzed the levels of uncleaved as well as polyadenylated snRNA transcripts, an indicative of 3’-end processing efficiency. For this purpose, endogenous INTS11 was knocked-down by co-transfecting an expression vector for dCAS9-KRAB with INTS11 promoter-specific guides (Figure 4A). This INTS11 depletion was then complemented either by the over-expression of a wild type GFP-FLAG-INTS11 (WT) or the different INTS11 mutants, from a heterologous promoter (53). Levels of endogenous and transfected GFP-FLAG-INTS11 proteins were assessed by western blot with an anti-INTS11 or anti GFP antibody (Figure 4A). Endogenous INTS11 was efficiently knocked-down as shown in figure 4A, leading to a clear increase in uncleaved (Figure 4B) and polyadenylated snRNAs (Figure 4C). As expected, transfected GFP-FLAG-INTS11 WT was able to restore basal levels of uncleaved and polyadenylated snRNAs, while the catalytically-inactive mutant INST11 E203Q was unable to do so. Remarkably, the SUMOylation deficient mutant, GFP-FLAG-INTS11 3KR, was also unable to rescue INTS11 depletion (Figure 4B and 4C). Consistently, similar results were obtained upon depletion of endogenous INTS11 by siRNA (Supplementary Figure S2A and S2B).

**Figure 4.**
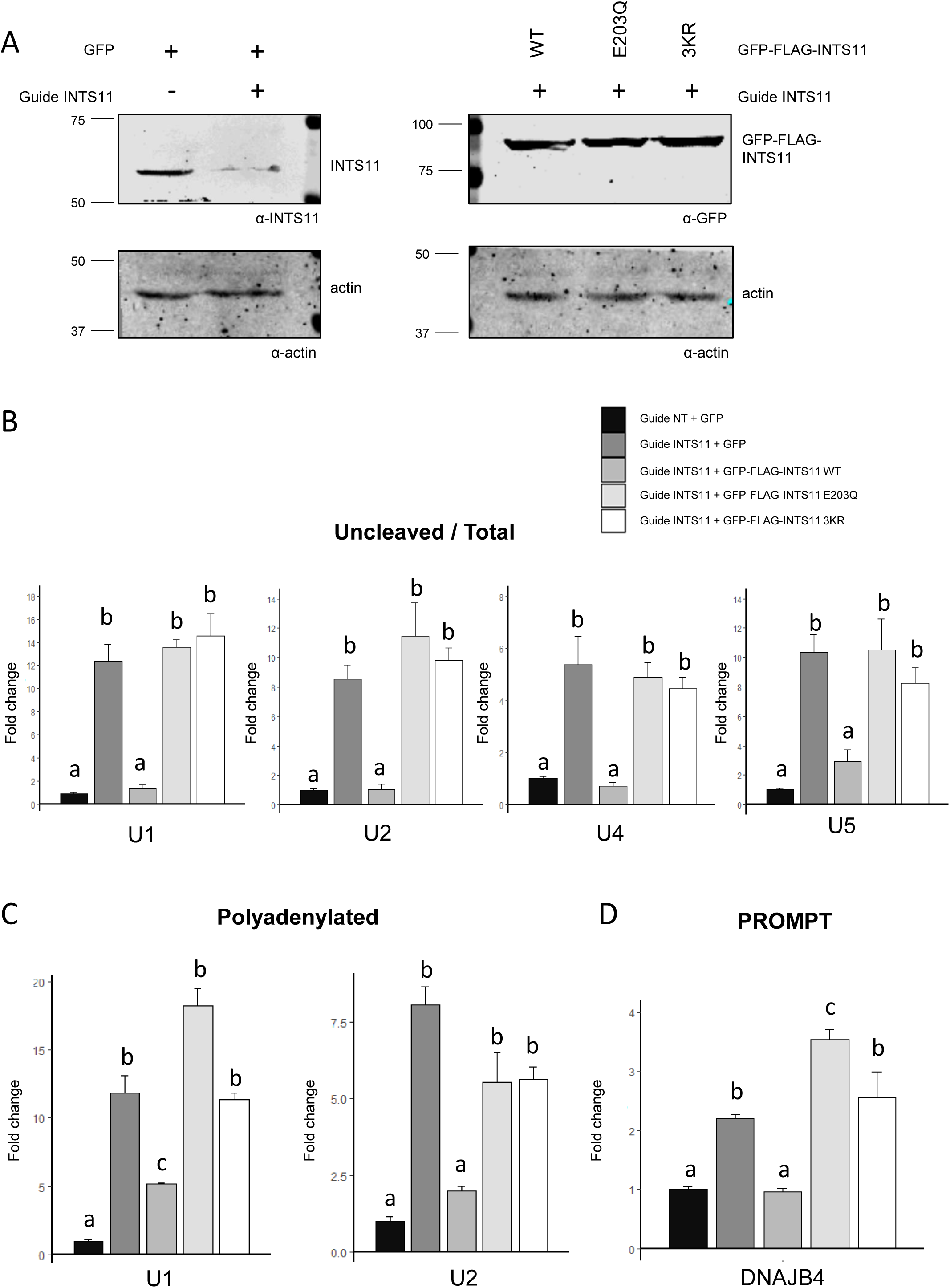
Lack of INTS11 SUMOylation affects Integrator activity in cultured cells. (A) HEK 293T cells were transfected with dCAS9-KRAB and a non-targeting guide (-) or a guide targeting INTS11 promoter (+) together with an expression vector for GFP (control), GFP-FLAG-INTS11 WT or GFP-FLAG-INTS11 mutated, as indicated above each lane. After 72 h, whole cell lysates were subjected to western blot with the antibodies indicated below each panel. (B-D) RT-qPCR analyses of RNA samples from cellular conditions indicated in A, using primer pairs targeting each of the indicated transcripts. Reverse transcription was performed with random decamer primer (B and D) or oligo dT primer (C). Transcripts levels were normalized to total snRNAs (B) or GAPDH mRNA (C-D). Average values with standard errors are shown. Bars labeled with the same letter are not statistically different (p > 0.05), while different letters indicate statistically significant differences. P-values were determined using a one-way ANOVA and Tukey post hoc test (n=4, p<0.001).

Over the last years, a growing body of experimental evidence has revealed that the Integrator complex also controls the processing and expression of other RNAPII-dependent transcripts beyond snRNAs, including enhancer RNAs (eRNAs), promoter upstream transcripts (PROMPTs), different long-noncoding RNAs (lncRNAs) and some mRNAs. We therefore selected several transcripts belonging to these different types of Integrator-regulated RNAs to further explore the involvement of SUMO in the diverse activities of the Integrator complex. To this end, we analyzed previously published TT-TimeLapse-sequencing data (TT-TL-seq) from INST11 depleted HEK 293T, rescued by the over-expression of INTS11 WT or the catalytically inactive mutant INTS11 E203Q (16) (Supplementary Figures S2C and S3A). In our hands, and in line with published TT-TL-seq data, the expression levels of the selected subset of transcripts increased upon depletion of endogenous INTS11. While over-expression of the WT version of this protein was able to restore transcripts basal levels, neither the E203Q nor the GFP-FLAG-INTS11 3KR mutants were able to do so (Figure 4D and Supplementary Figure S3B). These results are consistent with the idea that SUMOylation of INTS11 plays an important role for the function of the Integrator complex along its wide spectrum of RNA substrates.

### SUMOylation of INTS11 is relevant for its interaction with other Integrator subunits

To study a possible involvement of INTS11 SUMOylation in its interaction with protein partners, we co-transfected HEK 293T cells with expression vectors for GFP-FLAG-INTS11 or FLAG-INTS11, either WT or 3KR versions, plus different HA-tagged Integrator subunits. Whole cell lysates were subjected to co-immunoprecipitation (co-IP) using anti-GFP nanobodies coupled to magnetic beads or an anti-FLAG antibody. The co-precipitation of selected Integrator subunits was assayed via western blotting with anti HA antibody. INTS4 and INTS9, which associate with INTS11 forming the so-called “catalytic core” of the Integrator complex co-precipitate with INTS11 WT and INTS11 3KR to a similar extent (Figure 5A and 5B). The interaction of different INTS11 variants with INTS9 was also confirmed by co-expression of 6xHis-INTS9 and FLAG-tagged INTS11 followed by a nickel-mediated pull down of these two proteins (Supplementary Figure S3C). In contrast to what we observed with the catalytic module subunits, the levels of co-precipitated INTS13 and INTS14 with the INTS11 SUMOylation deficient mutant were drastically reduced in comparison to their co-precipitation with the WT version of INTS11 (Figure 5C and 5D). Taken together, these results suggest that SUMO conjugation to INTS11 is not necessary for its interaction with the other catalytic core components but is required for its interaction with other Integrator subunits, and consequently for proper assembly of this multimeric complex.

**Figure 5.**
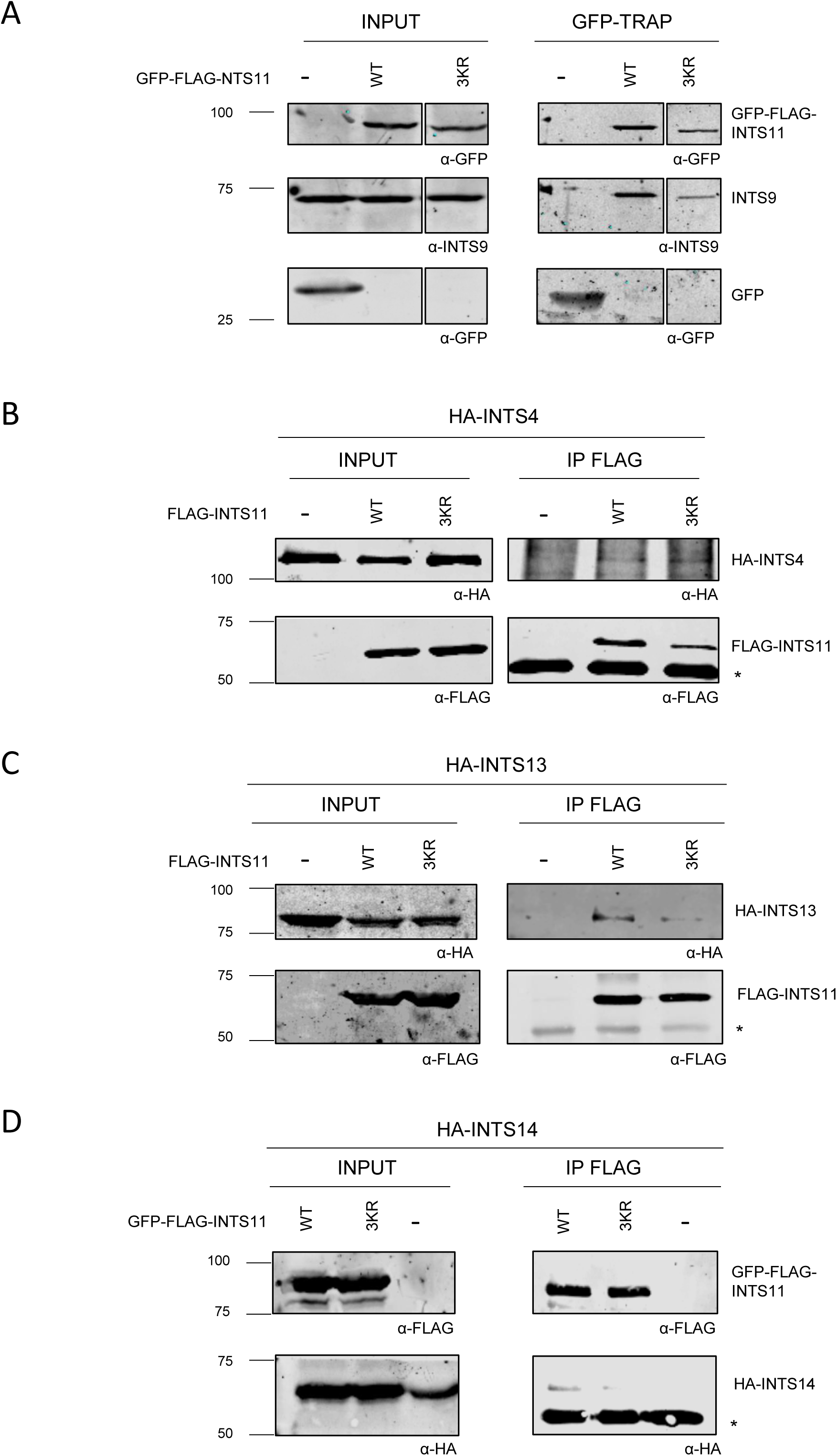
SUMOylation of INTS11 affects its interaction with other subunits of the Integrator complex. (A) Over-expressed GFP-FLAG-INTS11 WT or 3KR was immunoprecipitated from HEK 293T cell lysates using GFP-TRAP, and its association with INTS9 was assessed by western blot with the antibodies indicated below each panel. (B-D) Over-expressed GFP-FLAG-INTS11 WT or 3KR was immunoprecipitated with anti-FLAG from HEK 293T cell lysates containing HA-INTS4 (B), HA-INTS13 (C), or HA-INTS14 (D). Aliquots of whole cell lysates (input) and immunoprecipitates (IP) were analyzed by western blot with an anti-FLAG antibody to detect FLAG-INTS11 variants and anti-HA to detected enrichment of the other Integrator subunits. (* immunoglobulin heavy chain).

### INTS11 SUMOylation regulates Integrator sub-cellular localization

Considering the co-transcriptional nature of snRNA processing, we set out to determine whether SUMOylation affects INTS11 recruitment to chromatin. To this end, we performed a chromatin immunoprecipation (ChIP) assay with an anti-FLAG antibody and measured by qPCR the levels of precipitated 3’-box region corresponding to U2 gene. We observed significantly less association of the SUMOylation-deficient FLAG-INTS11 mutant to chromatin than of FLAG-INTS11 WT protein (Supplementary Figure S3D). Additionally, ChIP assays were performed using anti-INTS11 or anti-INTS9 antibodies, on extracts derived from INTS11-depleted cells either rescued with FLAG-INTS11 WT or with FLAG-INTS11 3KR. As expected, upon depletion of INTS11, less chromatin recruitment of this subunit was observed. In agreement with the functional heterodimerization of INTS11 and INTS9, less recruitment of INTS9 was also observed in INTS11-depleted cells (Figure 6A). Consistent with the results presented above, significantly less INTS9 and INTS11 associate with chromatin in the U2 3’-box region upon rescuing INTS11 depletion with FLAG-INTS11 3KR than with FLAG-INTS11 WT protein, even though similar expression levels of WT and 3KR FLAG-INTS11 were observed (Figure 6A, Supplementary Figure S4A and S4B).

**Figure 6.**
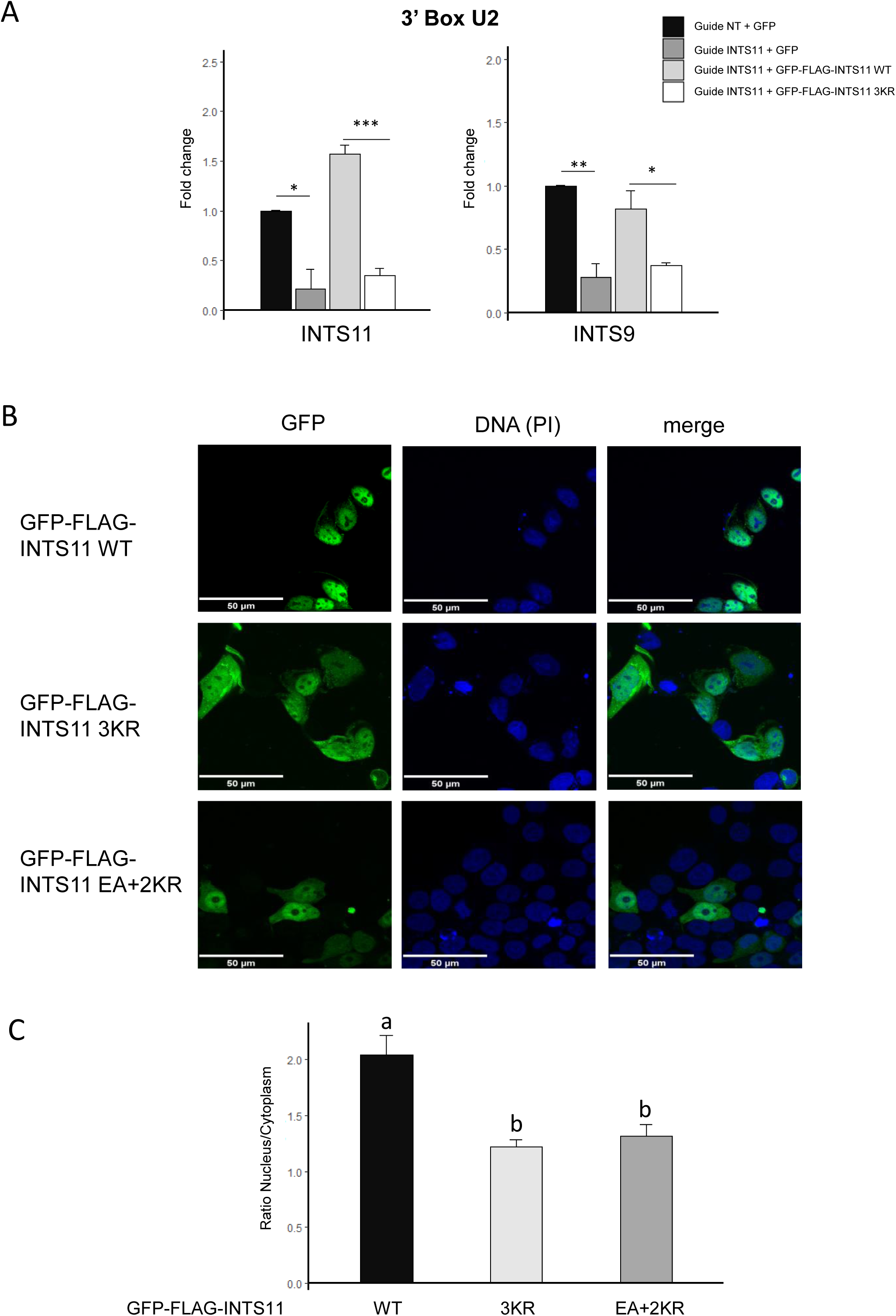
SUMOylation of INTS11 regulates its sub-cellular localization. (A) HEK 293T cells were co-transfected with dCAS9-KRAB and either a non-targeting guide (-) or a guide targeting INTS11 promoter (+), plus GFP, GFP-FLAG-INTS11 WT or GFP-FLAG-INTS11 3KR. After 72 h, ChIP analysis was performed with anti-INTS11 (left panel) or anti-INTS9 (right panel) antibodies. Quantification of immunoprecipitated DNA was assessed by qPCR with specific primers for U2 3’ box. Average values are shown with standard error (n = 3, ***P< 0.001; **P< 0.01; *P< 0.05; Student’s t test). (B) Representative confocal microscopy images of HeLa cells transfected with GFP-FLAG-INTS11 WT or GFP-FLAG-INTS11 mutated. Propidium iodide (PI)-nuclear staining is also shown. Scale bar, 50 µm. (C) Quantification of nucleus-cytoplasmic ratio from confocal microscopy images (n=150).

Previous studies have examined the sub-cellular localization of Integrator subunits with respect to CBs (54). It has been demonstrated that depletion of several Integrator subunit, including the catalytic core INTS4/INTS9/INTS11, causes the disassembly of CBs although these proteins do not show a robust colocalization with this sub-nuclear bodies (5).

Based on the results mentioned in preceding sections, we set out to analyze the possible impact of SUMO on INTS11 sub-cellular distribution. To do so, EGFP-INTS11 variants were over-expressed in HeLa cells and analyzed by fluorescence confocal microscopy. As already reported (5), EGFP-INTS11 WT displays a predominant nuclear localization (Figure 6B). Remarkably, EGFP-INTS11 3KR shows a perturbed localization as indicated by the decrease in its nuclear to cytoplasmic ratio (Figure 6B and 6C). Previous reports have demonstrated that the interaction of INTS11 with importins of the α type of nuclear transport receptor, as well as the nuclear localization signal (NLS) spanning from aa 460 to 479 within INTS11 were required for the nuclear localization of this protein (55).

To further explore whether INTS11 nucleocytoplasmic partitioning is regulated by SUMOylation, we first used bioinformatics prediction to search for putative NLS within INTS11 protein sequence. The software *NLS mapper* (http://nls-mapper.iab.keio.ac.jp/) recognizes the reported NLS as well as two others: a monopartite signal between aa 469 and 480 and a bipartite signal spanning from aa 565 to 597. We found that Lys 462, one of the residues that was replaced in our INTS11 3KR mutant, is strictly required for proper prediction of the bipartite NLS located at positions 460-479 (Supplementary Figure S4C). Based on this analysis, we generated a new INTS11 triple mutant termed “EA+2KR”. This INTS11 version keeps the Lys to Arg replacements at positions 381 and 475, as our previous SUMOylation deficient mutant. However, SUMOylation at Lys 462 is abolished not by the mutation of the target residue but instead, by altering the SUMOylation consensus motif (ΨKxD/E), replacing Glu 464 by Ala. Noticeable, this latter substitution does not affect the prediction of the reported NLS (460-479) (Supplementary Figure S4C). However, the subcellular localization of INTS11 EA+2KR is similar to that of INTS11 3KR (Figure 6B and 6C). In agreement with our previous observations, EA+2KR was unable to conjugate to SUMO (Supplementary Figure S4D) and to process 3’-end of snRNAs correctly (Supplementary Figure S4E and S4F). Taken together, these results indicate that INTS11 SUMOylation is necessary for its proper sub-cellular localization. In addition, considering that INTS11 3KR preserves its interaction with INTS4 and INTS9, at least in our co-IP analysis, the sub-cellular localization where the assembly of the catalytic core takes place represents an intriguing question.

## DISCUSSION

Despite the fact RNA-related proteins are the most abundant group among SUMOylation substrates (56–58), including many snRNA-related factors, very little is known about the regulation of proteins involved in snRNA biogenesis by SUMO conjugation. Previous studies have shown that depletion of the SUMO protease USPL1 causes a reduction in nascent and mature snRNAs levels, diminishes snRNPs production, and alters pre-mRNA splicing (51). However, these results could be linked to the relevance of USPL1 for assembly of CBs (50), sub-nuclear compartments where snRNA maturation occurs (1), which are in close proximity to where snRNA transcription takes place (48). To gain insight into the role of USPL1 in snRNAs biogenesis, we over-expressed this SUMO protease in cultured cells and found increased levels of uncleaved snRNAs (Figure 1), suggesting the balance between SUMOylation and de-SUMOylation is critical for snRNA maturation. This prompted us to investigate the impact of SUMO conjugation on the Integrator complex, which is responsible for snRNA 3’-end processing. Twelve out of fourteen subunits of this complex have been detected as SUMO conjugation targets by proteomic studies (27). However, it was not clear what consequences this PTM could have for Integrator complex activity. We confirmed that several of these Integrator subunits are *bona fide* SUMO substrates in cultured cells (Figure 2) and asked whether SUMO conjugation could be regulating the activity and/or the assembly of this complex. Although USPL1 knockdown does not alter global SUMOylation (50), we observe a slight reduction of global SUMO conjugation upon USPL1 over-expression (Figure 1D). Under this latter conditions, INTS11 SUMOylation levels are decreased but not those of other Integrator subunits tested (Figure 2I). Beyond this altered level of INTS11 SUMOylation, we cannot rule out that USPL1 could be regulating SUMO conjugation to different proteins involved in snRNAs biogenesis other than those tested in this work.

By site-specific mutagenesis, we obtained a mutant version of INTS11 that displays severely diminished SUMO conjugation levels, INTS11 3KR (Figure 3), which was further used along this study. INTS11 is responsible for 3’-end processing of snRNAs and associates with INTS4 and INTS9 to form the catalytic cleavage module (5). At first, we observed that INTS11 SUMOylation is necessary for efficient Integrator activity in 3’-end processing of snRNAs (Figure 4 and Supplementary Figure S3). Previous reports proposed additional roles for the Integrator complex within global transcription. It acts as an attenuator of expression for transcripts derived from weak promoters, like PROMPTS, eRNAs, and certain lncRNAs (7,11,12,14). However, in some cases, Integrator activity allows RNAPII to enter productive elongation and hence increases transcripcion (13,14). Here we show that INTS11 3KR is not capable of attenuating transcription (Figure 4D and Supplementary Figure S3B). Taken together, these results suggest that SUMOylation of INTS11 is necessary for its function not only in the context of snRNA maturation but also for regulating the expression of other RNAPII-dependent transcriptional units.

The formation of the INTS9/INTS11 heterodimer requires two specific domains in the C-terminal region of each of these proteins: CTD1 and CTD2 (5,6,52,59). It has been postulated that INTS9/INTS11 assembly requires an interaction between the CTD2 of INTS9 and the CTD2 of INTS11, followed by a gradual formation of multiple interactions between the CTD1 domains of both subunits. As a result, the dimerized CTD1s allow the recruitment of INTS4 to form the catalytic module (6). In this context, it has been shown that deletion of CTD1 from INTS11, where according to our work two SUMO acceptor residues are positioned (Figure 3D), does not impair the interaction between INTS9 and INTS11 (6). Consistent with these results, we observed that INTS11 3KR retains its interaction with INTS9 (Figure 5A and Supplementary Figure S3C). Furthermore, we show that the three mutations present within INTS11 3KR (K381/462R/475R) do not affect its interaction with INTS4 (Figure 5B). This indicates that SUMOylation of INTS11 is not required for the formation of the trimeric catalytic module and suggests that these mutations do not alter, at least considerably, the CTD1 domain. Recently, INTS10/INTS13/INTS14 has been characterized as an independent module, which binds to DNA and RNA and possibly stabilizes the cleavage module to target RNAs (60). We observed that INTS11 3KR displays a decreased interaction with two subunits of this module in comparison to the wild type version (Figure 5C and 5D). These results suggest that INTS11 SUMOylation is necessary to assemble the cleavage module into the whole Integrator complex.

The 3’-end processing of snRNAs occurs co-transcriptionally (61) and thus we performed ChIP analysis to assess whether the INTS11 SUMOylation-deficient mutant is recruited to chromatin as efficiently as the wild type version of this protein. The analysis reveals a diminished recruitment of INTS11 3KR to chromatin (Figure 6A and Supplementary Figure S3D). We further observed that INTS9 is less recruited to U2 gene loci in cells depleted of endogenous INTS11 than in cells with normal levels of this subunit, suggesting that INTS9 recruitment is dependent on INTS11. Consistent with the above-mentioned observation and with the fact that SUMOylation of INTS11 does not alter its interaction with INTS9, we detected less recruitment of INTS9 to chromatin when INTS11-depleted cells were rescued with INTS11 3KR, than when they were rescued with INTS11 WT (Figure 6A). Considering these results as well as the notion that SUMOylation regulates many cellular processes, including intracellular transport, we analyzed whether SUMO conjugation to INTS11 could alter its sub-cellular localization. By confocal microscopy, we show that GFP-FLAG-INTS11 3KR displays a decreased nuclear/cytoplasmic distribution ratio as compared to GFP-FLAG-INTS11 WT (Figure 6B and 6C). INTS11 possesses three putative NLSs predicted *in silico* by *cNLS mapper* (Supplementary Figure S4C). In particular, the signal encompassing residues 460 to 479 has been validated in cultured cells (55). By *in silico* analysis, we identified that the replacement of Lys 462 by Arg alters the score of the corresponding NLS, abrogating this signal, and thus possibly explaining the observed effect. To rule out that the observed change in sub-cellular localization is not merely due to the removal of the NLS by the single point mutation and to provide additional evidence for an involvement of SUMO conjugation in INTS11 sub-cellular localization, we generated an additional INTS11 mutant in which the SUMO consensus motif has been altered without affecting the NLS score (Supplementary Figure S4C). This mutant, which we called INTS11 EA+2KR, is unable to conjugate to SUMO and in line with our previous observations, fails to process nascent snRNAs (Supplementary Figure S4D, S4E and S4F). Remarkably, INTS11 EA+2KR display the same sub-cellular distribution that INTS11 3KR, pointing to the relevance of SUMO conjugation for proper INTS11 nucleus-cytoplasmic localization (Figure 6A and 6B). In addition, a previous study showed that Lys 462 is necessary for the correct 3’-end processing of snRNAs (6). Consistent with our results, this study reported that the mutation in this residue does not alter the formation of the catalytic module. Furthermore, they demonstrated that a highly positively charged composite tunnel formed by INTS9/INTS11/INTS4 is necessary for appropriate snRNA processing. These conclusions were achieved by altering the charges involved in this tunnel formation through mutating different residues in the participating proteins, including Lys 462 within INTS11 (6). Nevertheless, here we show that even without altering the charge of the 462 residue (replacing Lys by Arg), SUMOylation disruption in that position shifts the sub-cellular localization of INTS11. Thus, the results reported by Pfleiderer and Galej could also be interpreted in the context of the relevance of INTS11 SUMOylation for its proper sub-cellular distribution and consequently for its efficient activity.

It has been shown that INTS6/INTS8 form a module that recruit protein phosphatase 2 (PP2A) to active genes. This recruitment dephosphorylates CDK9 substrates, including RNAPII, and regulates the transcription cycle by pausing RNAPII (19,20). Furthermore, the structure of the Integrator complex bound to PP2A and paused RNAPII has been recently elucidated. This report helps to understand the mechanism by which the Integrator complex regulates transcription (62). However, almost nothing is known about the regulation of the function of this multimeric complex. Here, we demonstrate that INTS11 is SUMOylated and this post-translational modification is necessary to allow its proper sub-cellular localization, regulating the Integrator complex activity and possibly altering its assembly. The consequences of SUMO conjugation to other subunits of the complex remains to be explored.

## ACKNOWLEDGEMENTS

We thank Valeria Buggiano and Amaranta Avendaño for technical help; Shona Murphy, Joana Guiro, Maria Carmo-Fonseca, Joao Abreu Pessoa and other members from the Murphy and Fonseca laboratories for support, experimental suggestions and helpful discussion; Ezequiel Petrillo, Nicolas Nieto Moreno, and members of the Kornblihtt, de la Mata, Petrillo, Schor and Muñoz laboratories for valuable discussion; and Shona Murphy for critical reading of the manuscript.

