## Supplementary for "SUMO conjugation regulates the activity of the Integrator complex"

**Supplementary Material**

**SUPPLEMENTARY FIGURE LEGENDS**

**Supplementary Figure 1. INTS11 is SUMOylated at lysine residues 381, 462 and 475**. (A-B) HEK 293T cells were co-transfected with the expression vectors indicated at the top of each panel. After 48 h, whole cell lysates were subjected to nickel affinity chromatography (Ni-NTA). Aliquots of the cell lysates (input) and eluates (Ni-NTA) were analyzed by western blot with an anti-FLAG antibody to detect FLAG-INTS11 variants (A) or with an anti GFP antibody to detect GFP-FLAG-INTS11 (B). (C) Alignment of part of INTS11 protein sequence from human, rat, mouse, chicken, fruit fly and C. elegans showing the conservation of SUMO target sites and the corresponding consensus motifs.

**Supplementary Figure 2**. **Lack of INTS11 SUMOylation affects Integrator activity in cultured cells**. HEK 293T cells were co-transfected with control siRNA (-) or siRNA targeting INTS11 (+) and either an empty plasmid (-), or expression vectors encoding GFP-FLAG-INTS11 WT or GFP-FLAG-INTS11 mutated. After 72 h, whole cell lysates were subjected to western blot with an anti-INTS11 antibody or anti-tubulin. (B) RT-qPCR analyses of the RNA samples derived from the cultured cells indicated in A, using primer pairs targeting the indicated snRNAs. Reverse transcription was performed with oligo dT primer and transcripts levels were normalized to GAPDH mRNA. Bars labeled with the same letter are not statistically different ($p > 0.05$), while different letters indicate statistically significant differences. P-values were determined using a one-way ANOVA and Tukey post hoc test ($n=4$, $p<0.001$). (C) *Integrated Genomics Viewer* (IGV) genome browser screenshots corresponding to DNAJB4 from TT-TimeLapse sequencing (TT-TL- seq) experiments in HEK 293T cells, as published by Rosa-Mercado et al., 2021. Genes are oriented left to right. Scale is normalized to total mapped reads for each library.

**Supplementary Figure 3**. **Lack of INTS11 SUMOylation affects Integrator activity in cultured cells**. (A) *Integrated Genomics Viewer* (IGV) genome browser screenshots corresponding to PRPF38B and enhancer in chromosome 9 from TT-TL-seq experiments in HEK 293T cells, as reported by Rosa-Mercado et al., 2021. Genes are oriented left to right. Scale is normalized to total mapped reads for each library. (B) HEK 293T cells were transfected with dCAS9-KRAB and either a non-targeting guide (-) or a guide targeting INTS11 promoter (+), together with an expression vector for GFP, GFP-FLAG-INTS11 WT or GFP-FLAG-INTS11 mutated. After 72 h, RNA was subjected to RT-qPCR analyses using primers targeting the indicated transcripts. Transcript levels were normalized to GAPDH mRNA. Average values with standard errors are shown. Bars labeled with the same letter are not statistically different ($p > 0.05$), while different letters indicate statistically significant differences. P-values were determined using a one-way ANOVA and Tukey post hoc test (n=3, $p<0.001$). (C) HEK 293T cells were co-transfected with expression vectors for HIS-MYC-INTS9 and different FLAG-INTS11 variants, as indicated at the top of the panels. After 48 h, cell lysates were subjected to nickel pull-down. Aliquots of cell lysates (input) and eluates (pull-down) were analyzed by western blot with an anti-FLAG antibody to detect FLAG-INTS11 variants and anti-MYC antibody to detected enrichment of HIS-MYC-INTS9. (D) HEK 293T cells were transfected with either an empty plasmid (pcDNA) or expression vectors encoding FLAG-INTS11 WT or 3KR. After 48 h, ChIP analysis was performed with an anti-FLAG antibody. Quantification of immunoprecipitated DNA was assessed by qPCR with specific primers for U2 3' box. Data are represented as mean ± S.E. (n = 3, *$P < 0.05$; Student's t test).

**Supplementary Figure 4**. **SUMOylation of INTS11 regulates its sub-cellular localization**. (A-B) Whole cell lysates corresponding to cell culture conditions shown in Fig 6A were subjected to western blot with anti-INTS11, anti-FLAG, anti-INTS9 and anti-tubulin antibodies, as indicated at the bottom of each panel. (C) Score of INTS11 nuclear localization signals predicted by cNLS mapper. (D) HEK 293T cells were transfected with the expression vectors indicated at the top of each panel. After 48 h, cell lysates were subjected to nickel affinity chromatography (Ni-NTA). Aliquots of the cell lysates (Input) and eluates (Ni-NTA) were analyzed by western blot with an anti-GFP antibody to detect GFP-FLAG-INTS11 variants. (E) HEK 293T cells were transfected with dCAS9-KRAB and either a non-targeting guide (-) or a guide targeting INTS11 promoter (+), together with an expression vector for GFP, GFP-FLAG-INTS11 WT or GFP-FLAG-INTS11 mutated, as indicated above each lane. After 72 h, whole cell lysates were analyzed by western blot with anti-INTS11, anti-GFP and anti-actin antibodies. (F) RT-qPCR analyses of RNA samples corresponding to cell culture conditions shown in E, using primer pairs targeting the indicated snRNAs. Reverse transcription was performed with oligo dT primer. Transcripts levels were normalized to GAPDH mRNA.

**Supplementary Table S1**. Summary of Integrator proteins enriched by anti-SUMO2 immunoprecipitation. Data from Hendriks et al., 2018.

**Supplementary Table S2.** Primers used for RT-qPCR, qPCR, mutagenesis, and sequences of sgRNAs.

#### SUPPLEMENTARY MATERIALS AND METHODS

##### Transfection of siRNAs

siRNAs were transfected into HEK 293T and with Lipofectamine 2000 according to manufacturer's instructions (Thermo Fisher). The sequence corresponding to the human INTS11 3'UTR is UCUUUUGUUCAGCUUUUACUG and an siRNA targeting luciferase was used as a control, siRNA LUC: CUUACGCUGAGUACUUCGA(dT).

##### Nickel pulldown

HEK 293T cells were transfected in 35-mm culture wells with the indicated plasmids. After 48 h, cells were harvested and lysed in lysis buffer (50 mM Tris–HCl (pH 7.5), 500 mM NaCl, 10 mM Imidazol, 0,5 % (v/v) Triton) containing complete protease inhibitor (Roche). Extracts were sonicated at high amplitude with three 10-sec bursts, and insoluble material was pelleted. Ni-NTA beads (Qiagen) was added to the supernatant and incubated 1h with. Then, complexes were washed three times in wash buffer (50 mM Tris–HCl (pH 7.5), 500 mM NaCl, 30 mM Imidazol). For western blot analysis, samples were eluted in 2X Laemmli sample buffer containing 300 mM imidazole (pH 6.3) for 3 min at 95◦C

**Supplementary Figure S1**

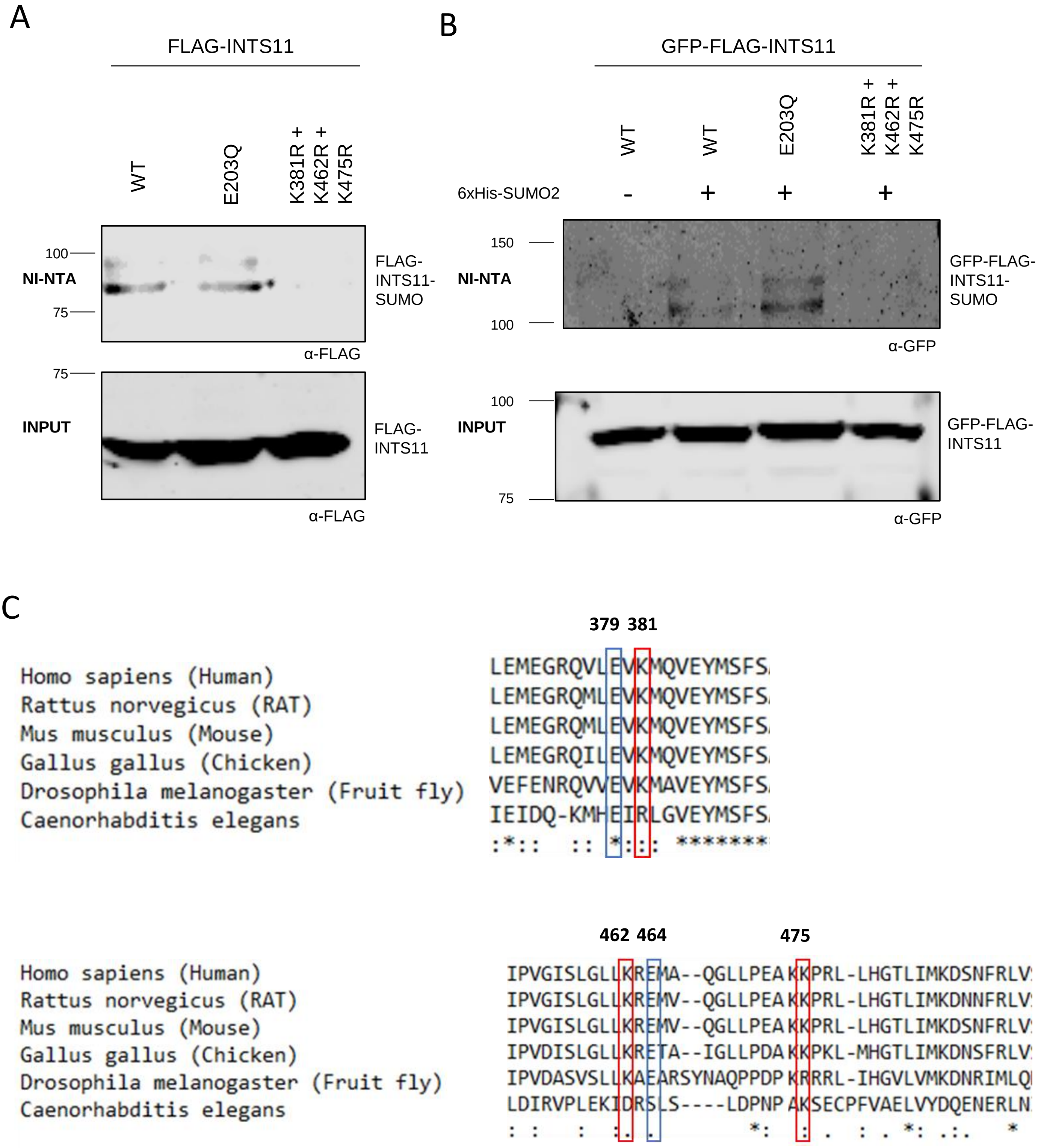

#### Supplementary Figure S2

A

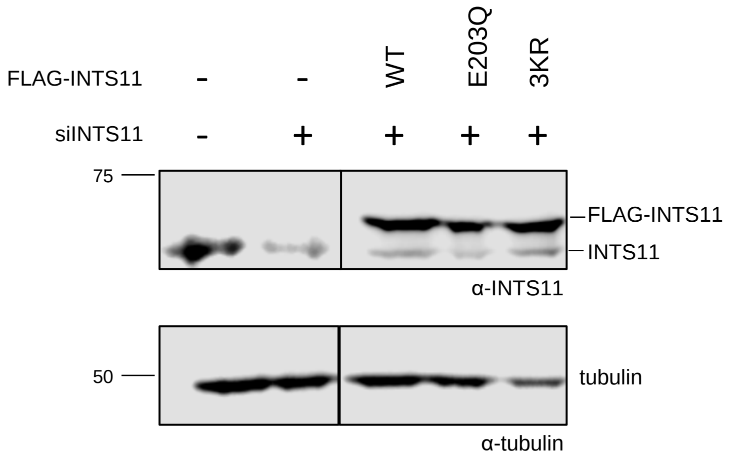


B

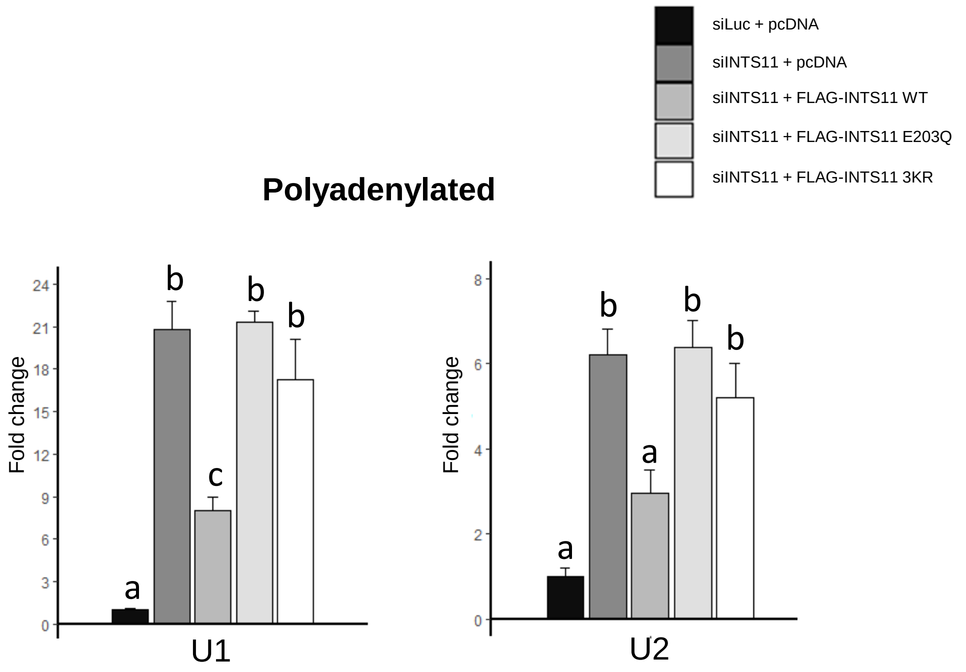


C

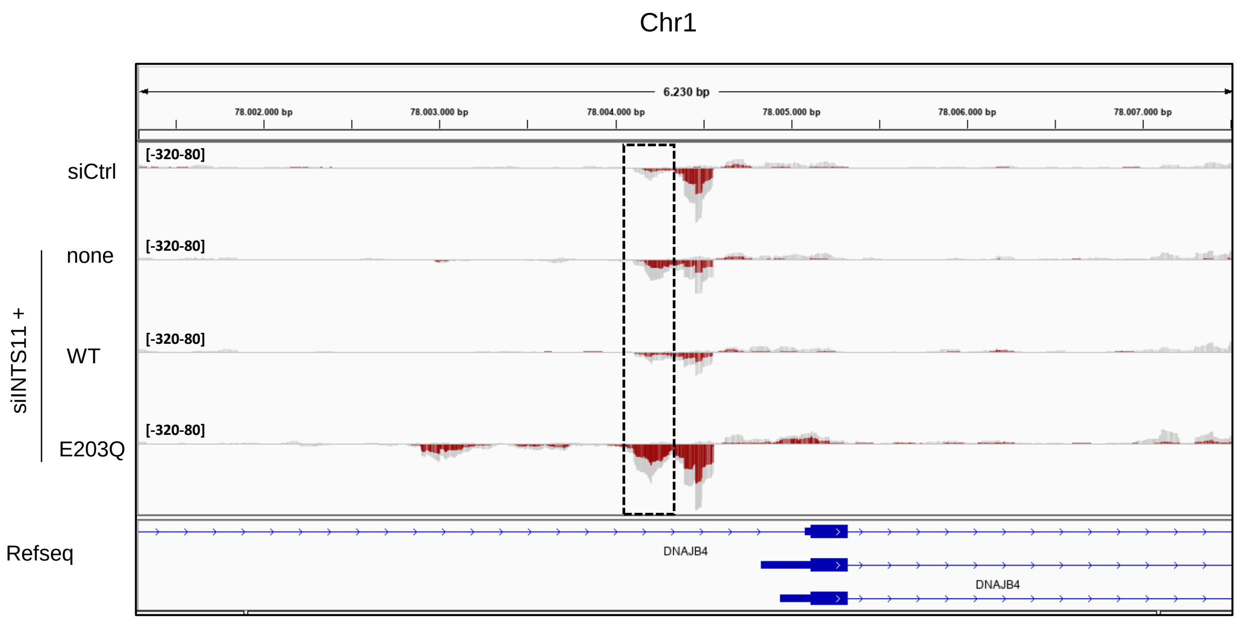

#### Supplementary Figure S3

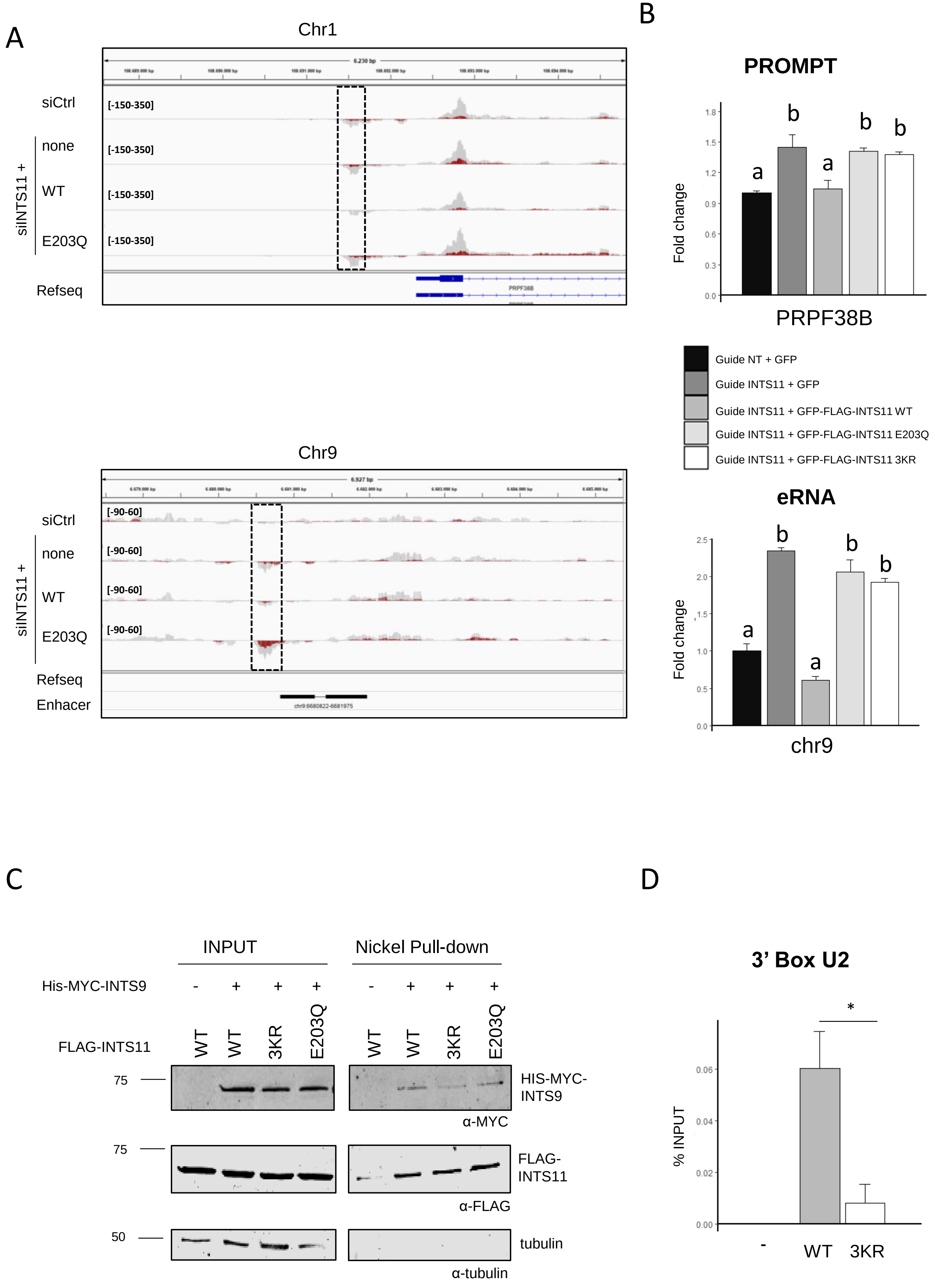

### Supplementary Figure S4

A

B

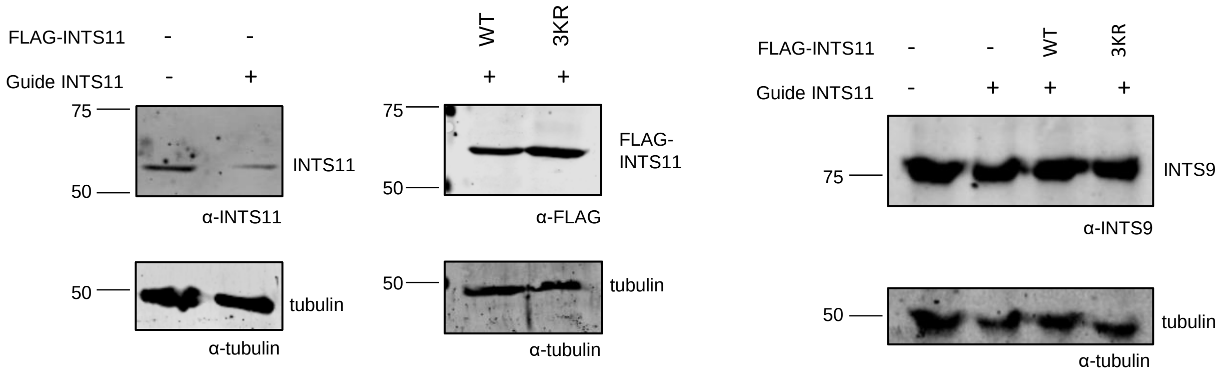


C

| Ints11 WT | | | |
|---|---|---|---|
| Pos | Sequence | NLS | Score |
| 469-480 | LLPEAKKPRLL | Monopartite | 8,5 |
| 460-479 | LLKREMAQGLLPEAKKPRL | Bipartite | 12 |
| 565-597 | EDPGTKVLLVSWTYQDEELGSFLTSLLKKGLP | Bipartite | 5,2 |

| Ints11 K381R | | | |
|---|---|---|---|
| Pos | Sequence | NLS | Score |
| 469-480 | LLPEAKKPRLL | Monopartite | 8,5 |
| 460-479 | LLKREMAQGLLPEAKKPRL | Bipartite | 12 |
| 565-597 | EDPGTKVLLVSWTYQDEELGSFLTSLLKKGLP | Bipartite | 5,2 |

| Ints11 3KR | | | |
|---|---|---|---|
| Pos | Sequence | NLS | Score |
| 469-480 | LLPEAKRPRLL | Monopartite | 10 |
| 460-479 | LLRREMAQGLLPEAKRPRL | Bipartite | ND |
| 565-597 | EDPGTKVLLVSWTYQDEELGSFLTSLLKKGLP | Bipartite | 5,2 |

| Ints11 K462R | | | |
|---|---|---|---|
| Pos | Sequence | NLS | Score |
| 469-480 | LLPEAKKPRLL | Monopartite | 8,5 |
| 460-479 | LLRREMAQGLLPEAKKPRL | Bipartite | ND |
| 565-597 | EDPGTKVLLVSWTYQDEELGSFLTSLLKKGLP | Bipartite | 5,2 |

| Ints E464A 2KR | | | |
|---|---|---|---|
| Pos | Sequence | NLS | Score |
| 469-480 | LLPEAKRPRLL | Monopartite | 10 |
| 460-479 | LLKRAMAQGLLPEAKRPRL | Bipartite | 15,3 |
| 565-597 | EDPGTKVLLVSWTYQDEELGSFLTSLLKKGLP | Bipartite | 5,2 |

| Ints11 K475R | | | |
|---|---|---|---|
| Pos | Sequence | NLS | Score |
| 469-480 | LLPEAKRPRLL | Monopartite | 10 |
| 460-479 | LLKREMAQGLLPEAKRPRL | Bipartite | 13,5 |
| 565-597 | EDPGTKVLLVSWTYQDEELGSFLTSLLKKGLP | Bipartite | 5,2 |

D

E

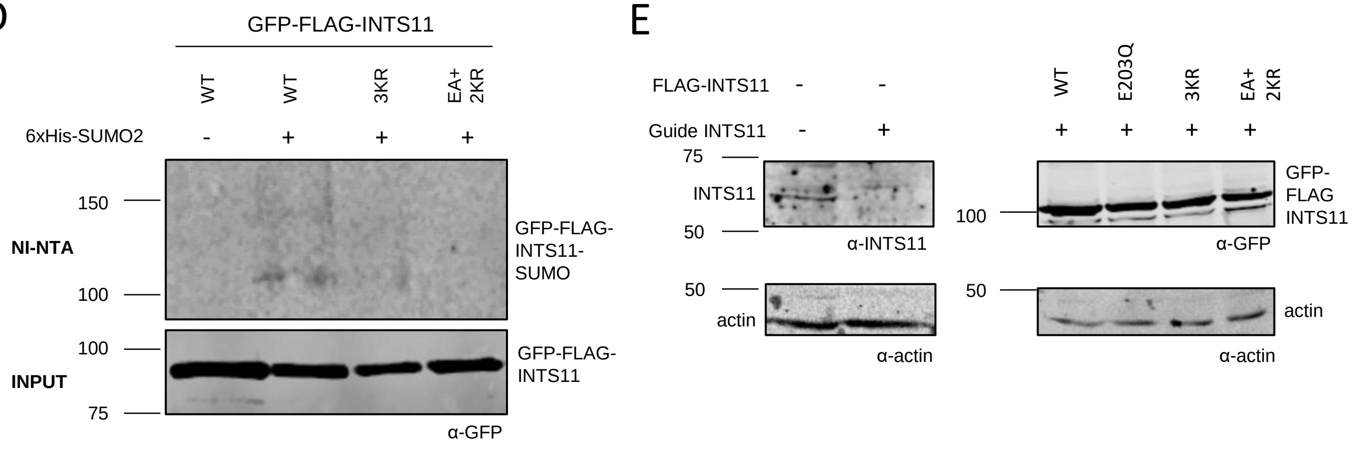


F

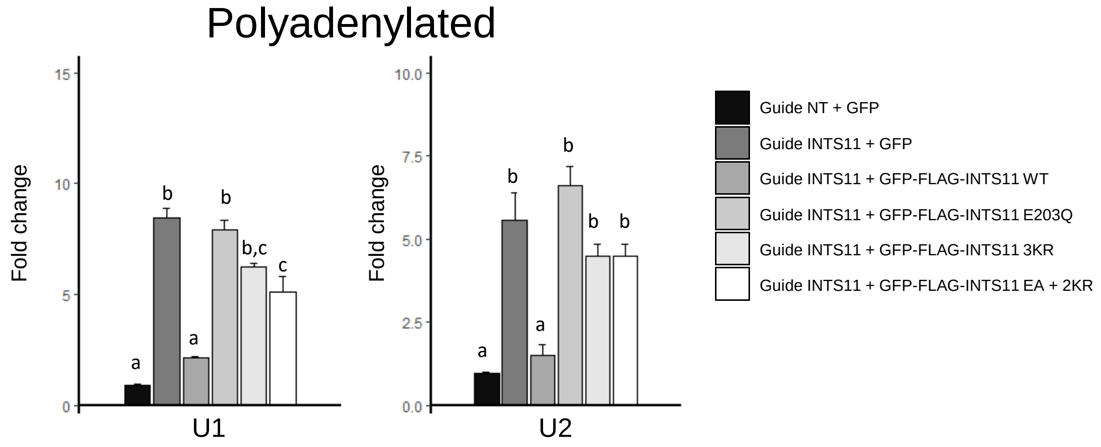

#### Supplementary Table S1

| Protein name | Gene name | Position |
|---|---|---|
| Integrator complex subunit 1 (Int1) | INTS1 | 33, 40, 47, 58, 76, 79, 100, 379, 524 |
| Integrator complex subunit 3 (Int3) | INTS3 | 480, 508, 549, 558, 986 |
| Integrator complex subunit 4 (Int4) | INTS4 | 15, 26, 169, 186, 198, 304, 324, 360, 593, 688, 791, 959 |
| Integrator complex subunit 6 (Int6) | INTS6 | 24, 275, 313, 451, 537, 559, 607, 611, 695 |
| Integrator complex subunit 7 (Int7) | INTS7 | 194, 277, 510, 785, 810, 847, 909 |
| Integrator complex subunit 8 (Int8) | INTS8 | 189, 244, 412, 436, 475, 708, 910 |
| Integrator complex subunit 9 (Int9) | INTS9 | 58, 170, 354, 459, 486, 515, 532, 550, 579, 582, 583 |
| Integrator complex subunit 10 (Int10) | INTS10 | 234, 239, 248, 400, 464, 539, 659 |
| Integrator complex subunit 11 (Int11) | INTS11 | 115, 234, 245, 251, 263, 274, 289, 316, 333, 367, 367, 369, 389, 426, 457, **462**, 471, **475** |
| Integrator complex subunit 12 (Int12) (PHD finger protein 22) | INTS12 | 15, 23, 58, 68, 72, 85, 105, 116, 117, 122, 225, 229, 242, 254, 309, 320, 329, 341, 347, 361, 370, 433, 452 |
| Integrator complex subunit 13 (Protein asunder homolog) | INTS13 | 2, 35, 167, 201, 224, 268, 295, 436, 507, 549, 589, 604, 611, 633, 648, 668, 690 |
| Integrator complex subunit 14 (von Willebrand factor A domain-containing protein 9) | INTS14 | 244, 340, 386, 406 |

### Supplementary Table S2

| Name | Sequence |
|---|---|
| **U1 snRNA Total FW** | CCATGATCACGAAGGTGGTTT |
| **U1 snRNA Total REV** | ATGCAGTCGAGTTTCCCACAT |
| **U2 snRNA Total FW** | TTCTCGGCCTTTTGGCTAAG |
| **U2 snRNA Total REV** | CTCCCTGCTCCAAAAATCCA |
| **U4 snRNA Total FW** | GCCAATGAGGTTTATCCGAGG |
| **U4 snRNA Total REV** | TCAAAAATTGCCAATGCCG |
| **U5 snRNA Total FW** | GGTTTCTCTTCAGATCGTATAAATC |
| **U5 snRNA Total REV** | CTCAAAAAATTGGTTTAAGACTCAGA |
| **GAPDH mRNA FW** | ATCGTGGAAGGACTCATGAC |
| **GAPDH mRNA REV** | GCAGGGATGATGTTCTGGAG |
| **U1 snRNAs Uncleaved FW** | TACCTGGCAGGGGAGATACC |
| **U1 snRNA Uncleaved REV** | GCGTACAGTCTACTTTTGAAACTC |
| **U2 snRNAs Uncleaved FW** | CTCCACGCATCGACCTGG |
| **U2 snRNA Uncleaved REV** | TGCTACCGTCTCTCACCTC |
| **U4 snRNAs Uncleaved FW** | CGTAGCCAATGAGGTCTATCCG |
| **U4 snRNA Uncleaved REV** | CTCTTCAACCTCCCAAAACC |
| **U5 snRNAs Uncleaved FW** | ATTTCCGTGGAGAGGAACAACTC |
| **U5 snRNA Uncleaved REV** | GCACCATTGAACAGAAAAAGGAA |
| **chr9 eRNA_1 FW** | GAGACCTGAGCTTACCCTCG |
| **chr9 eRNA_1 REV** | GGAGTTAAACTATGGAGCCGGTAC |
| **DNAJB4 prompt FW** | CGCAGGTTGTTTAAATTAGG |
| **DNAJB4 prompt REV** | CTCCCTTAACAATGTGAAAC |
| **Prpf38 B prompt FW** | CTTTTGTCGGGAGCTGAGCC |
| **Prpf38 B prompt REV** | CCAGGAGACCCCTCTCGGATAC |
| **INTS11 K115R FW** | CGCCGTAGACAAGAGGGGCGAGGCCAACTTC |
| **INTS11 K115R REV** | GAAGTTGGCCTCGCCCCTCTTGTCTACGGCG |
| **INTS11 FW K289R** | CTGGACCAACCAGAGGATCCGCAAGACTTTC |
| **INTS11 REV K289R** | GAAAGTCTTGCGGATCCTCTGGTTGGTCCAG |
| **INTS11 FW K369R** | CAGCGGGCAGCGGAGGCTCGAGATGGAGGG |
| **INTS11 REV K369R** | CCCTCCATCTCGAGCCTCCGCTGCCCGCTG |
| **INTS11 K381R FW** | GGTGCTGGAGGTCAGGATGCAGGTGGAG |
| **INTS11 K381R REV** | CTCCACCTGCATCCTGACCTCCAGCACC |
| **INTS11 K462R FW** | CGCTGGGGCTGCTGAGGCGGGAGATGGCGC |
| **INTS11 K462R REV** | GCGCCATCTCCCGCCTCAGCAGCCCCAGCG |
| **INTS11 FW K475R** | CCCTGAGGCCAAGAGGCCTCGGCTCCTGC |
| **INTS11 REV K475R** | GCAGGAGCCGAGGCCTCTTGGCCTCAGGG |
| **INTS11 E464A FW** | CTGCTGAAGCGGGCGATGGCGCAGGGGCT |
| **INTS11 E464A RV** | AGCCCCTGCGCCATCGCCCGCTTCAGCAG |
| **INTS11 E203Q FW** | AACCTGCTCATCACACAGTCCACGTACGCCA |
| **INTS11 E203Q REV** | TGGCGTACGTGGACTGTGTGATGAGCAGGTT |
| **FW guide NT** | GCAGGGTTTTCCCAGTCACGACGTTGTAAA |
| **REV guide NT** | CTTTACAACGTCGTGACTGGGAAAACCCTG |
| **FW guide INTS11 promoter** | CCACCCCACAGCCCTCCCGG |
| **REV guide INTS11 promoter** | CCGGGAGGGCTGTGGGGTGG |
